# Organizing Principles of Astrocytic Nanoarchitecture in the Mouse Cerebral Cortex

**DOI:** 10.1101/2021.11.05.467391

**Authors:** Christopher K. Salmon, Tabish A. Syed, J. Benjamin Kacerovsky, Nensi Alivodej, Alexandra L. Schober, Michael T. Pratte, Michael P. Rosen, Miranda Green, Adario DasGupta, Hojatollah Vali, Craig A. Mandato, Kaleem Siddiqi, Keith K. Murai

**Author notes:** Correspondence should be addressed to Dr. Kaleem Siddiqi or Dr. Keith Murai.

## Abstract

Astrocytes have complex roles in central nervous system (CNS) health and disease. Underlying these roles is an elaborate architecture based on frequent, extremely fine, but seemingly haphazard branches, as well as prominent features including tripartite synaptic complexes and perivascular endfeet. While broad categories of structures in astrocytes are known, the fundamental building blocks that compose them and their organizing principles have yet to be adequately defined. This is largely due to the absence of high-resolution datasets that can reveal nanoscopic features of astrocytes (i.e. 10-20nm diameter in x, y, and z) and a lack of computational approaches that can effectively interrogate astrocyte shape, organization, and nanoarchitecture. Here, we produced and analyzed multiple, high-resolution datasets of layer 2/3 mouse somatosensory cortex using focused ion beam scanning electron microscopy (8nm intervals) and computer vision approaches to provide a principled, quantitative analysis of astrocytic nanoarchitecture. A decomposition of astrocytes into fundamental ‘parts’ led to the discovery of unique structural components, recurring structural motifs, and assembly of parts into an organized hierarchy. New relationships were also discerned between astrocytic processes and other CNS microanatomy including mitochondria, tripartite synapses, and cerebrovasculature. By deploying computational resources to quantitatively understand the organizing principles and nanoarchitecture of astrocytes, this study reveals the specialized anatomical adaptations of these complex cells within the CNS.

**One Sentence Summary:** Using high-resolution serial electron microscopy datasets and computer vision, this study provides a systematic analysis of astrocytic nanoarchitecture from multiple samples of layer 2/3 of adult mouse neocortex, and presents quantitative evidence that astrocytes organize their morphology into purposeful, classifiable assemblies with unique structural and subcellular organelle adaptations related to their physiological functions.

## INTRODUCTION

Over a century ago, Ramon y Cajal and others proposed the Neuron Doctrine that seeded a coherent vision of connectivity, information flow, and capacity of the central nervous system (CNS) to carry out computations based on basic structural units –*axons, dendrites, and synapses*– of neurons (Ramón y Cajal, 1899). Yet astrocytes are the most common cells of the CNS, and they are now known to have diverse roles in CNS function, injury, and disease (Escartin et al., 2021; Haim and Rowitch, 2016; Khakh and Deneen, 2019). In the healthy CNS, they control ion/neurotransmitter homeostasis, synapse development/function, neurovascular coupling, metabolism, and maintenance of the blood brain barrier (BBB). While understanding information processing in the CNS by neurons is essential, this vision is incomplete without clarity on how astrocytes integrate signals from nearly every CNS cell type/structure (Santello et al., 2019) to participate in vital processes including respiration, vision, olfaction, food intake, memory recall, and sleep (Adamsky et al., 2018; Brancaccio et al., 2017; García-Cáceres et al., 2016; Halassa et al., 2009; Kim et al., 2014; Petzold et al., 2008)

A major challenge in deciphering astrocytic function is understanding their complex anatomical composition. This is a known bottleneck for the field (Khakh and Deneen, 2019; Rusakov, 2015) and one which limits our ability to attribute key astrocytic functions to their anatomy. The delicate features of astrocytes, comprised of myriad poorly defined branches that make up the majority of their structure, surpass the limit of detection using light and super-resolution microscopy (Khakh and Deneen, 2019). Protoplasmic astrocytes in gray matter are believed to have ‘sponge-like’ or spongiform anatomy and form mutually exclusive territories with neighboring astrocytes where their thousands of tiny processes interact with synapses (Bushong et al., 2002). Many of these processes are 10-20 nanometers in diameter, visualizable only by electron microscopy and require serial sectioning at thin interval sizes (Sandri et al., 1982; Spacek, 1985a; Ventura and Harris, 1999a). It is believed that the tips of these processes interact with pre- and postsynaptic sites through perisynaptic astrocytic processes (PAPs)(Papouin et al., 2017) at what is termed the tripartite synapse (Araque et al., 1999; Halassa et al., 2009; Perea et al., 2009). PAPs are sites where astrocytes locally detect synaptic activity and deliver neuroactive molecules such as ATP, glutamate, and D-serine to regulate synaptic transmission and plasticity (Araque et al., 2014). Astrocytes also form specialized endfoot compartments that encompass vasculature (Mathiisen et al., 2010; Simard et al., 2003). Endfeet exchange substances across the blood brain barrier (BBB), allow the CNS to adapt to metabolic load, and participate in remodeling of cerebrovascular tone (Gordon et al., 2007; Macvicar and Newman, 2015; Nuriya and Yasui, 2013). Despite the rapidly expanding knowledge of the importance of astrocytes in the CNS, an overall grasp of their nanoarchitecture remains incomplete. This hinders the ability to understand how astrocytes contribute to the connectome.

We built an experimental-computational pipeline to solve astrocytic nanoarchitecture in layer 2/3 mouse somatosensory cortex using high-resolution focused ion beam scanning electron microscopy (FIB-SEM) combined with computer vision analyses. This pipeline: (1) produced multiple datasets that sufficiently resolve the fine features of astrocytes (4.13 × 4.13 × 8nm voxel size), and (2) used computer vision to quantitatively analyze astrocyte geometry, complexity, and interactions with associated microanatomy. We uncovered a wealth of nanoscopic features related to astrocytic shape, organization, location/morphology of intracellular organelles, and interactions with other cell types/structures including synapses and vasculature. Contrary to what is believed, directed graphs representing layer 2/3 astrocytic architecture showed that astrocytes are not ‘sponge-like’ with frequent loops/holes. Instead, they are divisible into subcompartments organized hierarchically in a wide but shallow network. Shape analysis, unsupervised clustering, and 3-dimensional geodesic measurements show that astrocytes contribute diverse parts to tripartite synapses, have morphologically distinct populations of mitochondria, endoplasmic reticulum, and mitochondria-endoplasmic reticulum contact sites in their endfeet, display higher-order relationships with clusters of synapses, and strategically position mitochondria across their anatomy. Altogether, this study presents quantitative evidence of unique organizing principles in astrocyte structure and helps break the bottleneck for computational interrogation of astrocytic nanoarchitecture in the CNS.

## RESULTS

### Brain Samples, FIB-SEM, Feature Segmentation, and 3D Astrocyte Reconstructions

There are two main impediments to visualizing and analyzing astrocytic nanoarchitecture (Khakh and Deneen, 2019). <u>First</u>, astrocytes have an irregular anatomy composed of poorly defined branches that confocal and super-resolution light microscopy cannot fully resolve (Figure 1A-B)(Grosche et al., 1999; Khakh and Sofroniew, 2015). These frequent branches of varying thickness compose specialized subcompartments including tripartite synaptic complexes and endfeet. Many fine features of astrocytes are 10-20 nanometers in thickness, and are therefore most effectively visualized by serial electron microscopy (EM) with small slicing intervals (Sandri et al., 1982; Spacek, 1985a). <u>Second</u>, there is a dearth of computational tools and approaches capable of interrogating the structural complexity of astrocytes. Branching patterns of astrocytes have been described using surface-to-volume measurements on 60-70nm thick serial sections (Gavrilov et al., 2018; Patrushev et al., 2013). However, no studies have interrogated astrocytic geometry and topology at <10nm resolution (x, y, and z) that is required for proper resolution of fine astrocytic processes. New high-resolution datasets and computational methods are needed for a systematic, quantitative analysis of astrocytic nanoarchitecture in the healthy and diseased brain (Escartin et al., 2021; Khakh and Deneen, 2019).

**Figure 1.**
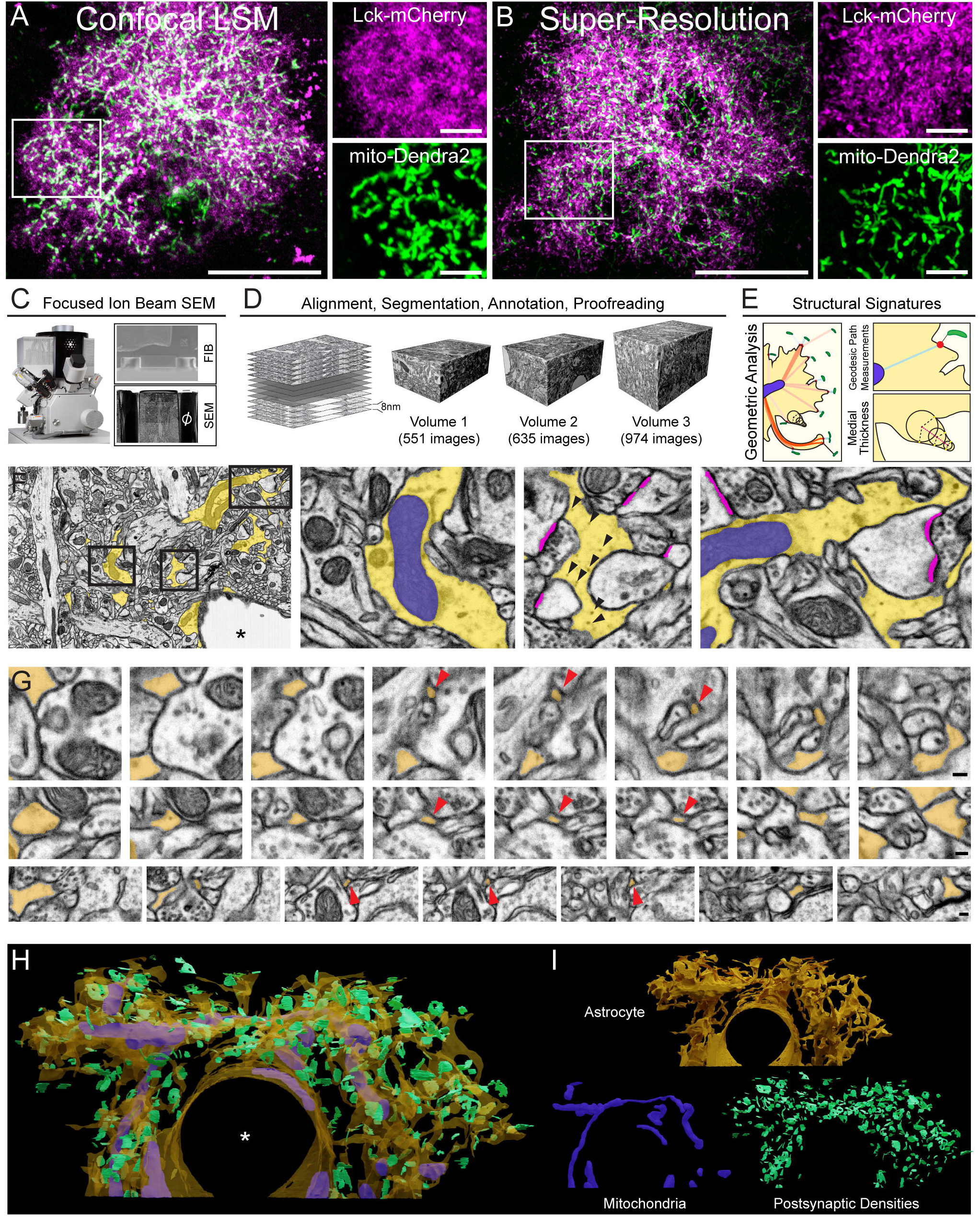
Astrocytes in adult mouse somatosensory cortex and astrocytic nanoarchitecture experimental-computational pipeline. (A) Astrocyte in somatosensory cortex visualized by confocal laser-scanning microscopy (LSM). In utero electroporation was performed at E18, delivering membrane-targeted mCherry (Lck-mCherry; magenta) and mitochondrial-associated Dendra2 (mito-Dendra2; green). (B) Super-resolution imaging by structured illumination microscopy reveals finer features of astrocytic anatomy than confocal microscopy but also has limitations in defining fine processes of astrocytes. (C) Serial electron microscopy (SEM) approach using focused ion beam scanning electron microscopy. Image to the left shows the instrumentation used (FEI Helios NanoLab 660 Dual Beam). Images to the right show typical sample configuration for focused ion beam (FIB) milling and SEM. (D) SEM stack processing, alignment, feature segmentation, annotation, and proofreading. Three references volumes from Layer 2/3 S1 cortex used in the study are shown. (E) Geometric analysis involving geodesic path and medial thickness measurements. (F) Example SEMs with astrocyte (yellow), mitochondria (blue), and PSD (magenta) segmentation. Arrowheads point to electron dense glycogen granules in the relatively translucent astrocyte cytoplasm used to identify astrocytic processes. (G) Example SEM series’ showing the nanoscopic size of curving astrocytic structures. Red arrowheads point to fine features of astrocytes that are resolvable with small sectioning intervals (32-40nm interval shown). (H) 3D reconstruction of the astrocyte in Volume 2 (yellow; Astro 2; 635 serial images, 4.13nm axial resolution) with mitochondria (blue), a large capillary endfoot, and contacting postsynaptic densities of excitatory synapses (green). Scale = 20um (low magnification) and 4um (high magnification) in A and B; 100nm in G. Calibration cube = 1μm.

To address these challenges, we developed an experimental-computational pipeline for multi-sample quantitative analysis of astrocytic nanoarchitecture (Figure 1C-E). We utilized focused ion beam scanning electron microscopy (FIB-SEM) to produce three uninterrupted ultrastructural series’ of layer 2/3 of 7-week old mouse somatosensory (S1) cortex (mill depth of 8nm and axial resolution of ∼4.13 nm)(Figure 1C-D). At 10,000x magnification and 3,072 × 2,048 resolution, fine nanoscopic processes of astrocytes were clearly resolved in reference volumes, while providing a wide field of view to image large parenchymal and perivascular subcompartments (Figure 1F). All volumes contained neuropil that allowed direct comparisons between reference volumes. An 8nm FIB sectioning interval was selected in order to detect fine astrocytic processes and to avoid split, merge, and connectivity errors known to occur with astrocytic processes at 29nm sectioning intervals (Kasthuri et al., 2015).

We performed alignment of FIB-SEM image stacks and segmentation of astrocytic volumes using TrakEM2 (Cardona et al., 2012). Astrocytes were identified in electron micrographs by their relatively translucent cytoplasm, absence of neurofilaments, and presence of electron-dense glycogen granules (Figure 1F). We initially screened each volume for thick profiles of astrocytes giving rise to smaller branches that extended extensively into the volume. Next, six individuals focused on manual segmentation of astrocytic volumes and associated structures and two individuals performed quality control assessments. Selection of the 8nm section interval proved to be critical, as many very fine branches (<20nm) of the astrocytic membrane were observed (Figure 1G). The segmentation process yielded multiple, high-quality reconstructions of astrocyte sub-volumes and prominent features of interest: mitochondria, endoplasmic reticulum (ER), and postsynaptic densities (Figure 1H-I). All reference volumes, segmented datasets and reconstructions from this study will be made available to the scientific community for further analysis.

### Nanostructural Parts and Signatures of Layer 2/3 Astrocytes in Mouse S1 Cortex

Currently available software/algorithms are under-powered to deal with computational problems associated with astrocytic nanostructure such as their complex geometry and topology, and the sheer number of intricate processes. We therefore designed computer vision approaches to define and quantify these specific structural attributes. For nanoarchitectural analysis we performed the following: (i) Tips of processes were identified by searching for surface points that were spatially distant from the average loci of other points in their local spatial neighborhood (see Methods). This allowed us to define a measure of “protrusiveness” for all points on the astrocytic surface (Figure 2A). (ii) We computed the medial surface within the volume using average outward flux-based skeletonization algorithms. (iii) We then mapped the radius of the medial sphere centered at each point of the medial surface back to the boundary to yield a measure of local thickness for the astrocyte processes (Figure 2B)(Siddiqi and Pizer, 2008). (iv) Finally, by combining protrusiveness and local thickness measurements, we obtained a single continuous structural measure (M-score), representing a range of surface point characterizations, from narrow tips to thick core regions. When qualitatively exploring the complex 3D structure of the segmented portions of astrocytes, we observed that astrocytic processes have distinct thick and thin regions, and that the processes do not simply become progressively thinner toward their tips. We therefore used the M-score to classify key part types composing the astrocyte: thin ‘constrictions’ present both at branch tips and internally within the branching astrocytic structure; thicker ‘expansions’; and the thickest trunk- or pillar-like ‘core’ regions (Figure 2C-D). The most identifiable and basic structural motif we found upon examining the reconstructions, aside from the abundance of constrictions at the ends of processes and scarcity of core regions, were expansions which were separated from one another by narrow constrictions (Figure 2E). A comparison of astrocytes from three independent reference volumes showed consistency in the proportion of astrocytic part types (Figure 2F). This consistency across astrocytes was also found when comparing surface areas, volumes, and surface area-to-volume ratios (S/V) of constrictions and expansions (Figure 2G). The same was true when comparing thickness and protrusiveness measures (Figure 2H). Astrocyte 3 showed some slight differences in mean thickness (thicker cores, thinner expansions and constrictions)(Figure 2H). This is likely related to Astrocyte 3 being a larger, more complete sample of astrocyte processes and also being fully embedded in neuropil rather than near other structures such as a blood vessel for Astrocyte 2.

**Figure 2.**
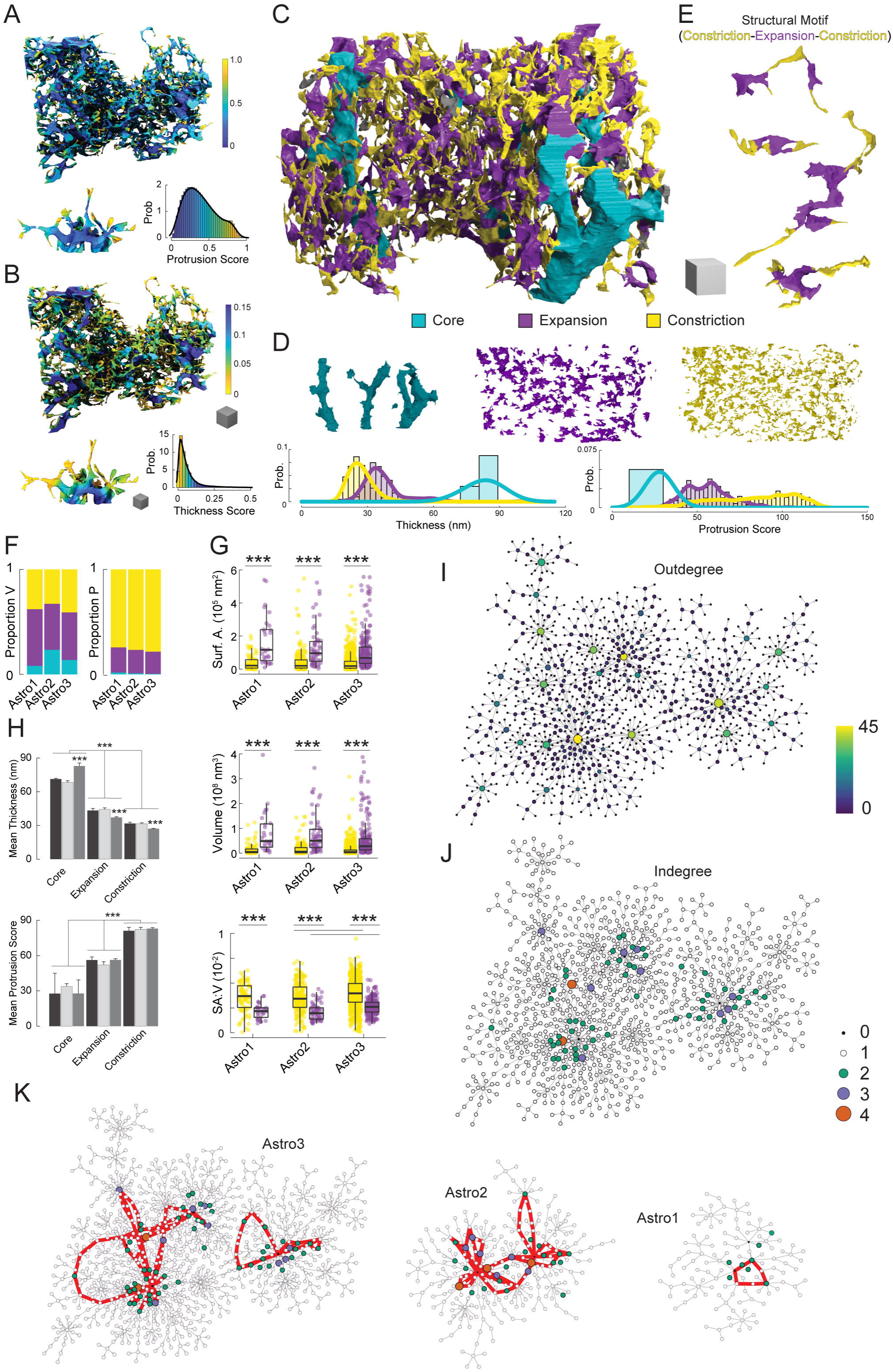
Astrocytic nanoarchitectural analysis. (A) Protrusion scores mapped onto Astro 3 with a heatmap designating structures from high (yellow) to low (blue) protrusiveness. (B) Thickness scores mapped onto Astro 3 a heatmap designating structures from high (blue) to low (yellow) thickness. (C) Identification of astrocytic parts (cores, expansions and constrictions) by combining protrusion and thickness scores (M score). (D) Top, gallery view of astrocytic cores, expansions, and constrictions. Note the scarcity of cores and abundance of expansions and constrictions. Bottom, probability density distribution of mesh surface vertices based on thickness (in nm) or protrusion score. (E) Identification of the commonly found constriction-expansion-constriction structural motif. (F) Comparison of the proportion of volume and parts (cores, expansions, and constrictions) from astrocytes from 3 references volumes. (G) Comparison of expansions and constrictions based on surface area, volume, and surface-to-volume ratio for astrocytes in 3 reference volumes. (I) Graph of outdegree of Astro 3, revealing the relatively shallow, but wide network of subcompartments. Only a few parts give rise to large connectivity, i.e., parts/hubs (vertices) with up to 45 connections (edges). (J) Graph of indegree to identify where looping structures occur (i.e. where indegree is greater than 1). (K) Following inspection of 3D models, few looping structures are found in astrocytes from three reference volumes. *** p < 0.001

It is difficult to appreciate the complex organization of astrocytes solely through visualization of 3D reconstructions. Thus, in order to understand the potential organization of astrocytic structure (i.e. whether sponge-like or strictly hierarchical), we interpreted the cores, expansions, and constrictions as separate nodes in a graph. Starting at the core regions, edges were placed in a direction away from cores towards their connecting parts, from which we proceeded in a recursive manner to construct further edges to additional connected parts, until a terminal part was reached. The resulting directed acyclic graph allowed us to analyze the hierarchical nature of astrocytic parts. Graphing outdegree revealed a relatively shallow, but wide network of astrocytic parts, with only a few parts giving rise to large connectivity, i.e., hub parts with up to 45 outward edges (Figure 2I). Large hubs eventually led to a dispersed population of intermediate-size hubs, which were still a few edge lengths away from the terminal nodes. Historically, astrocytes have been widely believed to have a ‘sponge-like’ or spongiform anatomy with a perforated architecture containing abundant holes and loops, and with presumably random organization (Arizono et al., 2020; Bushong et al., 2004; Savtchenko et al., 2018; Schiweck et al., 2018; Verisokin et al., 2021). This interpretation is largely based on impressions from light microscopy or lower resolution EM visualization of astrocytes and not founded on quantification of astrocytic shape in 3D space. To achieve clarity on this, we examined node indegree in 3 reference volumes, to pinpoint where looping structures occurred. Loops in directed acyclic graphs must contain at least one node with indegree greater than 1, indicating the presence of more than one incoming path at that node (Figure 2J). Following inspection of 3D models in which parts contributing to putative looping structures were examined in isolation, we found that looping structures are present in L2/3 astrocytes (Figure 2K). However, loops were relatively scarce which is uncharacteristic of true sponge morphology (Chen et al., 2017). Rather, astrocytes mainly consisted of independent branches of undulating thickness, emanating from core regions and ending at terminal parts without looping back on themselves.

### Structural Basis of the Tripartite Synapse

The classical description of the tripartite synapse suggests that PAPs occur at the tips of astrocytic processes that contact pre- and/or postsynaptic sites with most occurring near spines and postsynaptic densities (PSDs)(Figure 3A)(Arizono et al., 2020; Panatier et al., 2014). This subcellular compartment is believed to be equipped for local detection of synaptic activity and release of gliotransmitters to regulate synaptic transmission and plasticity (Araque et al., 2014). Tripartite synapses have been characterized in previous ultrastructural studies (Genoud et al., 2006; Ventura and Harris, 1999b; Witcher et al., 2007, 2010). However, while their neuronal pre- and post-synaptic components are readily identifiable, little is known about what unique astrocytic features actually constitute the tripartite synapse. Given the high resolution of our FIB-SEM datasets and ability to account for every synapse in contact with the astrocytic surfaces examined, we directly addressed this question. We focused on asymmetric postsynaptic densities (PSDs) of spine synapses, which define the postsynaptic sites of excitatory synapses and whose size correlates with synaptic efficacy (Murthy et al., 2001; Ostroff et al., 2002; Schikorski and Stevens, 1997). We segmented all PSDs in contact with astrocytes in three reference volumes. This yielded 2,107 segmented PSDs in total. Volumes of PSDs fell into a right-skewed Rayleigh-like distribution as observed previously (Figure 3B)(Bartol et al., 2015). Average PSD volumes were similar between reference volumes 1 and 2, with PSD volumes smaller for reference volume 3 (Figure 3B). However, anatomical characteristics of segmented PSDs were similar between the three reference volumes, with respect to synapse type (asymmetric/asymmetric), location (shaft/spine), presence of a spine apparatus, and perforation (macular/perforated)(Figure 3C)(Mishchenko et al., 2010; Spacek, 1985b). Dendritic spines and asymmetric synapses accounted for the vast majority of synapses. A subset of spines contained a spine apparatus, the Ca^2+^- modulating extension of smooth ER into spines (Breit et al., 2018), and a mostly overlapping but distinct subset of spines also possessed a perforated PSD, which have been seen to increase in number following stimuli inducing synaptic potentiation (Figure 3C)(Nikonenko et al., 2002; Sorra et al., 1998). As expected, the presence of spine apparatuses and perforations correlated with larger PSDs (Figure 3D)(Spacek and Harris, 1997; Yuste and Bonhoeffer, 2001). Combined, these measurements suggest that we examined astrocytes contacting synapses that have expected properties and distribution.

**Figure 3.**
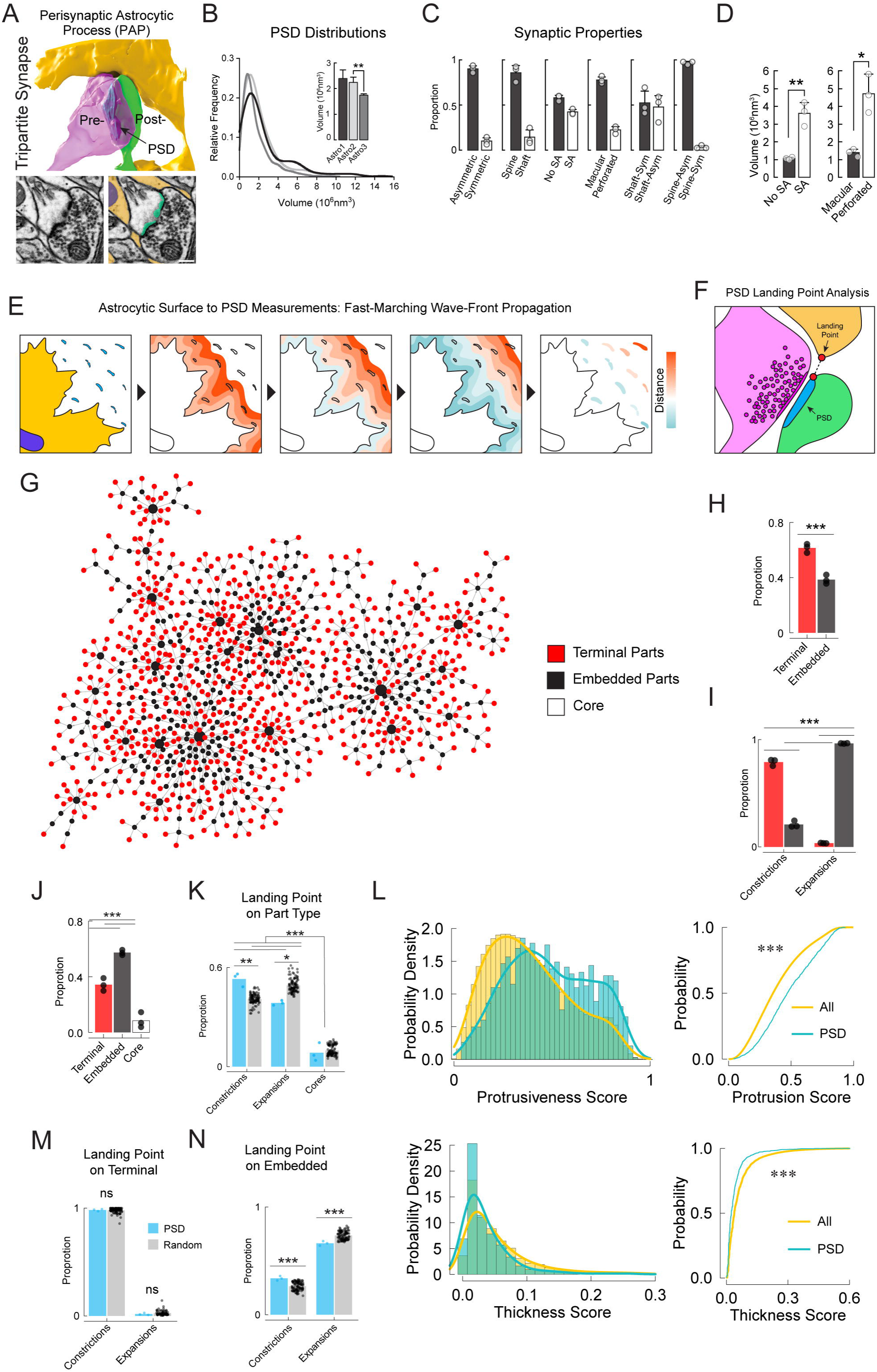
Tripartite synapse analysis. (A) An example reconstructed tripartite synapse from a reference volume. A dendritic spine is shown in green with a PSD shaded darker (arrow). An axonal bouton is shown in magenta. (B) Distribution of PSD volumes showing right-skewed Rayleigh-like distributions across three reference volumes. (C) Comparison of the properties of synapses in three references volumes. (D) PSDs associated with spine apparatus’ and perforated architecture show larger volume. (E) Geodesic path measurement approach to identify synaptic/PSD landing points on the astrocytic surface. A sequence depicting wave-front evolution from the surface of the astrocyte. (F) Schematic depicting the tripartite synapse. A dendritic spine is shown in green with its PSD shaded darker. An axonal bouton is shown in magenta with its active zone and vesicles shaded darker. An astrocyte is depicted in yellow with a PSD landing point indicated (arrow). (G) Left, graph of Astro 3 showing terminal (red), embedded (black), or core (white) astrocytic parts. (H) Bar graph showing that the proportion of parts that are terminal are significantly greater than the proportion of embedded parts across three reference volumes. (I) The proportion of constrictions and expansions that are either terminal or embedded in three reference volumes. (J) The proportion of PSD landing points that are terminal, embedded, or on core parts. (K) The proportion of actual PSD landing points versus randomly assigned landing points based on available surface vertices. Constrictions have an over-representation of landing points while expansions have an under-representation of landing points. (L) Most PSD landing points occur on terminal constrictions. (M) Both expansions and constrictions receive PSD landing points. Scale bar = 200nm. * p<0.05, ** p<0.01, *** p<0.001

Having verified synaptic populations within our reference volumes, we then computationally analyzed tripartite synapses across three reference volumes. Using 3D geodesic path measurements, we identified synaptic/PSD “landing points” on the astrocytic surface and cross-referenced these points against astrocytic nanostructure (Figure 3E-F). Given the classical view of the tripartite synapse being composed by thin PAPs that are at the ends of astrocytic processes, it might be predicted that the majority of tripartite synapses involve constrictions. On the other hand, as astrocytes detect and respond to synaptic activity with Ca^2+^ signals and gliotransmitter release, the larger size of astrocytic expansions might afford astrocytes the ability to utilize nearby organelles and other structures to perform biochemical processes at the tripartite synapse. Core regions would be the least expected astrocytic part to contain a PAP and contribute to tripartite synapses considering their thickness and large size. We first determined the proportion of astrocytic parts that could be classified as terminal, embedded, or core within astrocytic nanoarchitecture (Figure 3G). We utilized node degree to separate terminal parts (indegree = 1, outdegree = 0) from embedded parts (outdegree ≥ 1, or indegree >1 in the case of parts creating a loop). This analysis revealed a significant majority of astrocytic parts are terminal parts located at the ends of processes, as opposed to embedded parts that are flanked on either end by other parts (terminal, 61.5 ± 1.9%; embedded, 38.5 ± 1.9%, p = 0.0009, n=3 astrocytes)(Figure 3H). Samples contained only 2 to 4 cores of widely varying volume and thus cores were omitted form this analysis. Refining this comparison, we found that constrictions were mainly at process terminals but still composed an appreciable number of embedded parts, whereas expansions where almost exclusively embedded (Figure 3I).

Synapses have been shown to be evenly distributed within cortical layers (Merchan-Perez et al., 2014). Therefore, given the uneven distribution of astrocytic part type (and thus part volume and surface area), we wondered whether specific part types preferentially constitute tripartite synapses. Interestingly, PSD landing points were significantly more likely to be found on embedded parts (Figure 3J), despite the preponderance of terminal parts as a percentage of total parts (Figure 3H). This is likely due to the fact that embedded parts account for almost all expansions, which contain more vertices, surface area and volume (Figure 2F-G). To account for this volume factor and investigate the possibility that the location of PSD landing points (i.e. tripartite synapses) is not entirely random, we compared the landing points to sets of randomly placed points on the astrocyte surface (33 sets of points per astrocyte, each set equal in number to the PSD landing points for that astrocyte). This revealed a significant overrepresentation of landing points on constrictions and underrepresentation on expansions (Figure 3K). However, a surprising number of landing points were found across the entire distributions of protrusiveness/thickness scores (Figure 3L) and part types (Figure 3K), suggesting that PAPs, while having a strong representation on constrictions, are not located there obligately.

We next asked what part types were favored for landing points on terminal vs embedded parts. Nearly all landing points on terminal parts occurred on constrictions. This was also true for randomly seeded points on terminals (Figure 3M). Deeper inside the astrocytic territory, *en-passant* landing points favored expansions over constrictions (Figure 3N). This can be associated with the larger volume proportion and size of expansions (Figure 2F-G). However, it should be noted that *en-passant* contacts of synapses with constrictions occur more often than expected (Figure 3N). Thus, while the astrocytic component of the tripartite synapse is often a terminal constriction, many tripartite synapses also involve compartments embedded deeper within the hierarchy of astrocytic nanostructures, creating *en-passant* tripartite synapses on both constrictions and expansions.

### Nanoscopic Analysis of Astrocytic Mitochondria, Endoplasmic Reticulum (ER), and Mitochondrial-ER Contact Sites (MERCs)

The shape of mitochondria has been linked to their bioenergetic properties and provides a mechanism through which mitochondria balance ATP production according to local energy demands and nutrient supply (Liesa and Shirihai, 2013a; Mishra and Chan, 2016; Sebastián and Zorzano, 2018; Tondera et al., 2009). While it is known that mitochondria in protoplasmic astrocytes occupy about 6.5% of the perinuclear cytoplasmic volume, a detailed analysis of astrocytic mitochondrial shape and the interactions of astrocytic mitochondria with other subcellular organelles such as ER remain to be better understood. To investigate mitochondrial ultrastructure in astrocytes, we divided our astrocytic models into two broad regions with distinct roles, perivascular endfeet and the ‘non-endfoot’ regions that display a higher degree of morphological complexity and which we previously had subdivided into constituent parts. To bolster the analysis, we added a fourth reference volume that contained a large, cross-sectional area of a capillary with endfeet (Figure 4; Supplementary Figure 1). Comparison of reconstructed mitochondria from endfoot and non-endfoot regions revealed significant differences in the shape of mitochondria in these astrocytic subcompartments. While non-endfoot mitochondria were largely composed of long and short tubules, mitochondria in endfeet displayed a flattened, often disk-like, morphology (Figure 4A). This was reflected in quantitative analysis of mitochondria in the endfoot which showed lower surface area-to-volume ratios (S/V)(Figure 4B; p = 0.0011), lower average surface curvatures (Figure 4C; p = 0.007), and higher average thicknesses (Figure 4D; p = 0.037) than mitochondria in non-endfoot regions. We also observed that endfoot mitochondria were often thinned towards the centre, with a concave surface (Figure 4E). The specialized morphology of mitochondria in endfeet may contribute to metabolic adaptations of this specialized subcompartment involved in gliovascular coupling.

**Figure 4.**
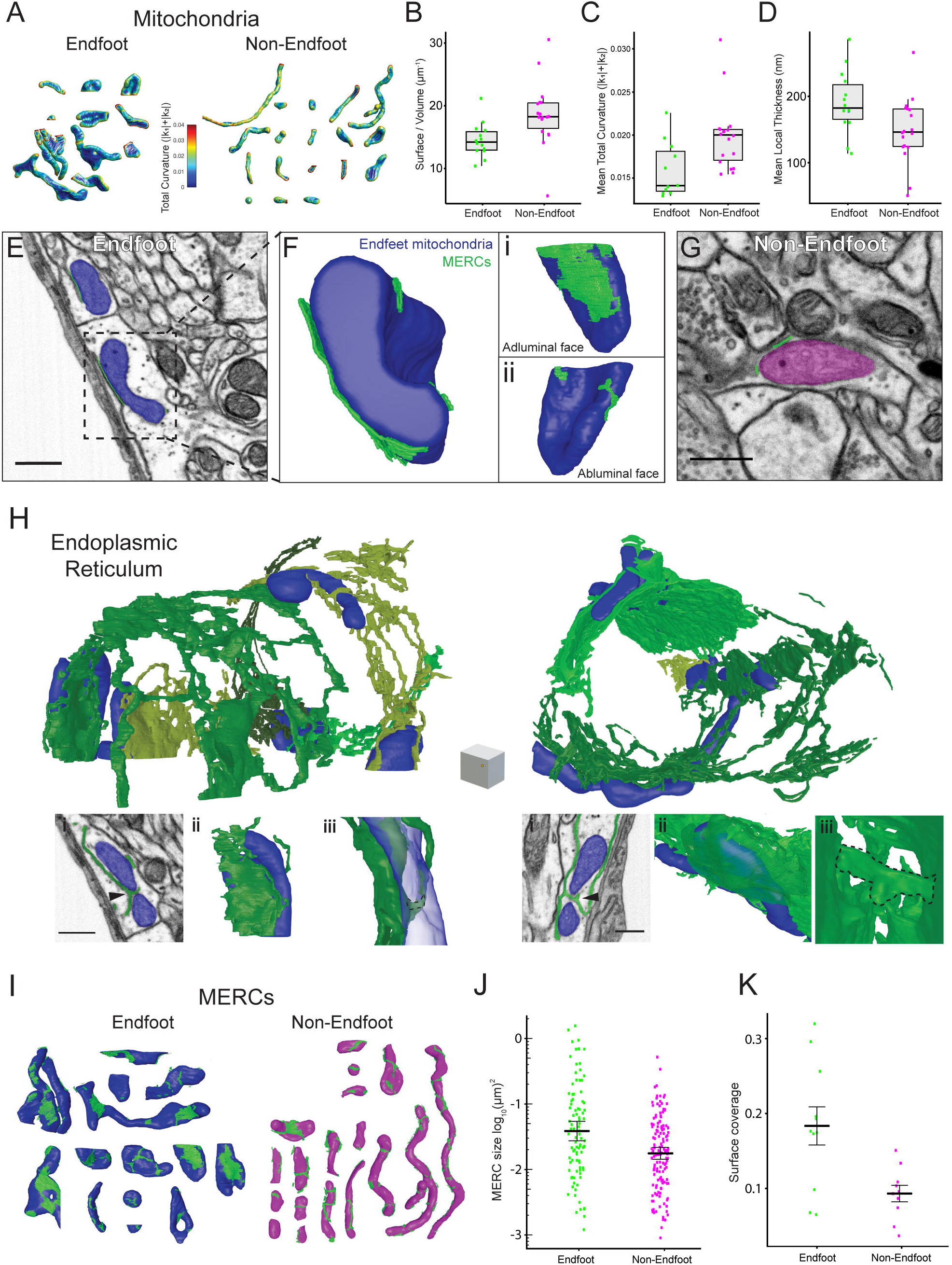
Nanostructural analysis of astrocytic mitochondria, endoplasmic reticulum (ER), and mitochondria-ER contact sites (MERCs). (A) Reconstructed astrocytic mitochondria from two reference volumes in endfeet (left) and neuropil-associated parts (right). Surface colour indicates total curvature (|k1|+|k2|). (B-D) Surface-to-volume ratio (B; p = 0.0011; Mann-Whitney test; n = 14 endfoot, 18 non-endfoot mitochondria from 2 reference volumes), mean total surface curvature (C; p = 0.007; Mann-Whitney test; n = 14 endfoot, 18 non-endfoot mitochondria from 2 reference volumes), and mean local thickness (D; p = 0.037; Mann-Whitney test; n = 14 endfoot, 18 non-endfoot mitochondria from 2 reference volumes) of mitochondria in endfeet and neuropil-associated parts. (E-F) SEM of endfoot mitochondria and MERCs. MERCs were often found to be particularly prominent on the side of mitochondria which faced the lumen of blood vessels (adluminal surface). (G) SEM of mitochondria in neuropil-associated parts. (H) Two separate reconstructions of endfoot ER. ER was typically observed to be closely associated with the flattened portions of mitochondria (i-iii) and in some cases ER was seen to pass through the centre of “doughnut-shaped” mitochondria (i, arrowheads; iii, dotted area). (I) Reconstructed mitochondria in endfeet (blue) and neuropil-associated processes (magenta) with reconstructed MERCs (green). (J-K) Quantitative comparison of individual MERC size (J; p = 0.0016; Welch’s unequal variance t-test on log transformed data; n = 85 endfoot, 106 non-endfoot mitochondria from 2 reference volumes) and the fraction of mitochondrial surface occupied by MERCs (K; p = 0.006; Welch’s unequal variances t-test; n = 14 endfoot, 10 non-endfoot mitochondria from 2 reference volume) between astrocyte compartments. Scale = 400nm in SEM in E, G, and H. Scale cube = 500 × 500 × 500 nm^3^ in H.

Mitochondria work in tandem with the endoplasmic reticulum (ER) to fulfill several critical roles in intracellular Ca^2+^ signaling and homeostasis (Patergnani et al., 2011). To study astrocytic ER, we reconstructed large parts of the ER network in FIB-SEM reference volumes (Figure 4E-H). We found that most of the ER in non-endfoot regions consisted of a polygonal array of tubules connected by three-way junctions as previously described (Shibata et al., 2006)(Supplemental Figure 1), with occasional small sheet-like regions. In contrast, ER reconstructed in endfoot subcompartments showed a distinct organization with a large portion of the ER forming large, sheet-like cisterns (Figure 4H). To quantify the relative amounts of ER tubules and cisterns, we calculated the surface curvature of the reconstructed ER segments since this is a key distinguishing feature between the two types of ER structures observed (Supplemental Figure 1A)(Shemesh et al., 2014; Shibata et al., 2006; Westrate et al., 2015). We found that the distribution of curvature values could be well-approximated using a Gaussian mixture model with three components. Based on this model we assigned points on the surface of the ER into three clusters and mapped them onto the surface mesh of the reconstructed ER (Supplemental Figure 1B-C). Cluster 1 contained ER with the lowest curvature, and described the flat, sheet-like membranes of ER cisterns. Cluster 2 corresponded to tubular networks and the rounded edges of sheets (Shemesh et al., 2014). Cluster 3 revealed small surface protrusions with high curvature (Supplemental Figure 1B). In astrocytic non-endfoot regions, ∼88% of ER surface was found in Cluster 2, reflecting its highly tubular structure. Only 3%-6% of ER in non-endfoot regions fell into Cluster 1. Remarkably, ER in the endfoot displayed a different structural composition, with a larger proportion of points attributed to Cluster 1 (12%-39% in reference volumes)(Supplemental Figure 1B). Close examination of endfoot ER cisterns that account for this shift toward lower curvature also revealed a large number of fenestrations (holes in the sheet)(Shemesh et al., 2014; Terasaki et al., 2013), lined by cluster 2-type ER tubules (Supplemental Figure 1D).

Another unique feature of ER cisterns in astrocytic endfeet was their arrangement in two layers, with one layer wrapping around the capillary in close apposition to the adluminal astrocytic cell membrane. In many instances, we found mitochondria nestled between two sheets, forming extensive interaction sites with the surrounding ER (Figure 4H). The lack of helicoidal membrane “ramps” connecting the two sheets suggested that these membrane arrangements were structurally distinct from the membrane stacks commonly found in cell bodies (Shemesh et al., 2014; Terasaki et al., 2013). Notably, we found that ER in endfeet were often associated with the flat portions of mitochondria, and, in two cases we observed ER tubules passing through the centre of flat mitochondria highlighting the tight interaction between the two organelles in the endfoot (Figure 4H).

Cooperative interactions between mitochondria and ER occur at specialized sites called mitochondria-ER contacts sites (MERCs). MERCs are important sites of Ca^2+^ exchange between mitochondria and ER. They further allow the two organelles to cooperate in lipid and protein biosynthesis, and are involved in disease states (Marchi et al., 2017; Raturi and Simmen, 2013; Simmen et al., 2010). Because of their functional importance, we reconstructed MERCs from two FIB-SEM reference volumes to investigate structural differences between MERCs in endfeet and non-endfeet regions. MERCs were defined as sections of ER membrane running in parallel to the mitochondrial membrane, with a cleft of 30nm or less (Giacomello and Pellegrini, 2016), and within range for detection using an 8nm slicing interval. The electron dense nature of MERCs, which is likely related to a high concentration of tethering and signalling proteins, also served as a visual cue for identifying and tracing these contact sites. A commonly used metric for comparing the extent of MERC formation is the amount of mitochondrial outer membrane involved in contact sites (Hirabayashi et al., 2017)(Figure 4I). Individual reconstructed MERCS in endfeet were significantly larger compared those in non-endfeet regions (Figure 4J; endfeet 0.038µm^2^, parts 0.018µm^2^; p = 0.0016). Surprisingly, we found that the average MERC coverage of endfeet mitochondria was nearly double that of non-endfeet mitochondria (Figure 4K, endfeet 18.3+/-2.5%; parts 9.3+/-1.1%; p = 0.006). These results show that astrocytes have specific structural adaptations of their mitochondria, ER, and MERCS, and that these structures vary significantly between endfeet and non-endfeet compartments.

### The Structural Properties of Excitatory Synapses Do Not Influence Their Proximity to the Astrocytic Surface or Mitochondria

Given the importance of the interaction between astrocytic structure, synapses, and mitochondrial function/trafficking (Agarwal et al., 2017; Derouiche et al., 2015; Gӧbel et al., 2020; Jackson and Robinson, 2018; Stephen et al., 2015), we developed computational approaches to understand their inter-relationship by measuring 3D distances inside and outside astrocytic nanostructure (Figure 5A). Previously, distance measurements in serial electron microscopic reconstructions have been performed manually or indirectly along surfaces, centerline approximations and radii of intersecting spheres (Jorstad et al., 2014, 2018; Patrushev et al., 2013). We therefore designed an approach that would allow for accurate and scalable measurements of distances between points within the convoluted morphology of astrocytes. Based on observations that mitochondria give rise to a significant portion of Ca^2+^ events in astrocytes (Agarwal et al., 2017) and that astrocytic mitochondria display activity-related mobility related to glutamate transporter activity (Jackson et al., 2014), we hypothesized that mitochondrial position in astrocytes is influenced by the structural properties of nearby synapses (Figure 5A). To test this, we measured distances between all PSDs and their nearest mitochondria (PSD-Mito distances). To do so, we first measured distances from PSDs to their nearest point on the astrocyte surface (PSD landing points, Figure 3). We then performed a second round of eikonal equation-based distance measurements in which we evolved a 3D wave-front from the mitochondria to the astrocytic surface (Astro-Mito)(Figure 5B). This allowed for a shortest path to be measured from any point on the surface of the astrocyte to the nearest mitochondrion. The sum of pairs of PSD-Astro (Figure 3F) and Astro-Mito distances gave PSD-Mito distances (Figure 5C). PSD-Mito distances varied in astrocytes across the three reference volumes (Figure 5D). We observed that synapses varied widely in their proximity to astrocytic mitochondria, from <1um and up to more than 25um (Figure 5D,E). Despite this broad range, PSD volume was not correlated with proximity to mitochondria across three datasets (Figure 5E). Furthermore, no PSD attributes (i.e. asymmetric vs. symmetric; spine vs. shaft, presence or absence of spine apparatus; macular or perforated) were associated with either shorter or longer distances to their nearest mitochondrion (Figure 5F). These results suggest that PSDs predicted to be associated with stronger synapses (i.e. larger, perforated PSDs with spine apparatuses) do not have closer spatial relationships with astrocytic mitochondria.

**Figure 5.**
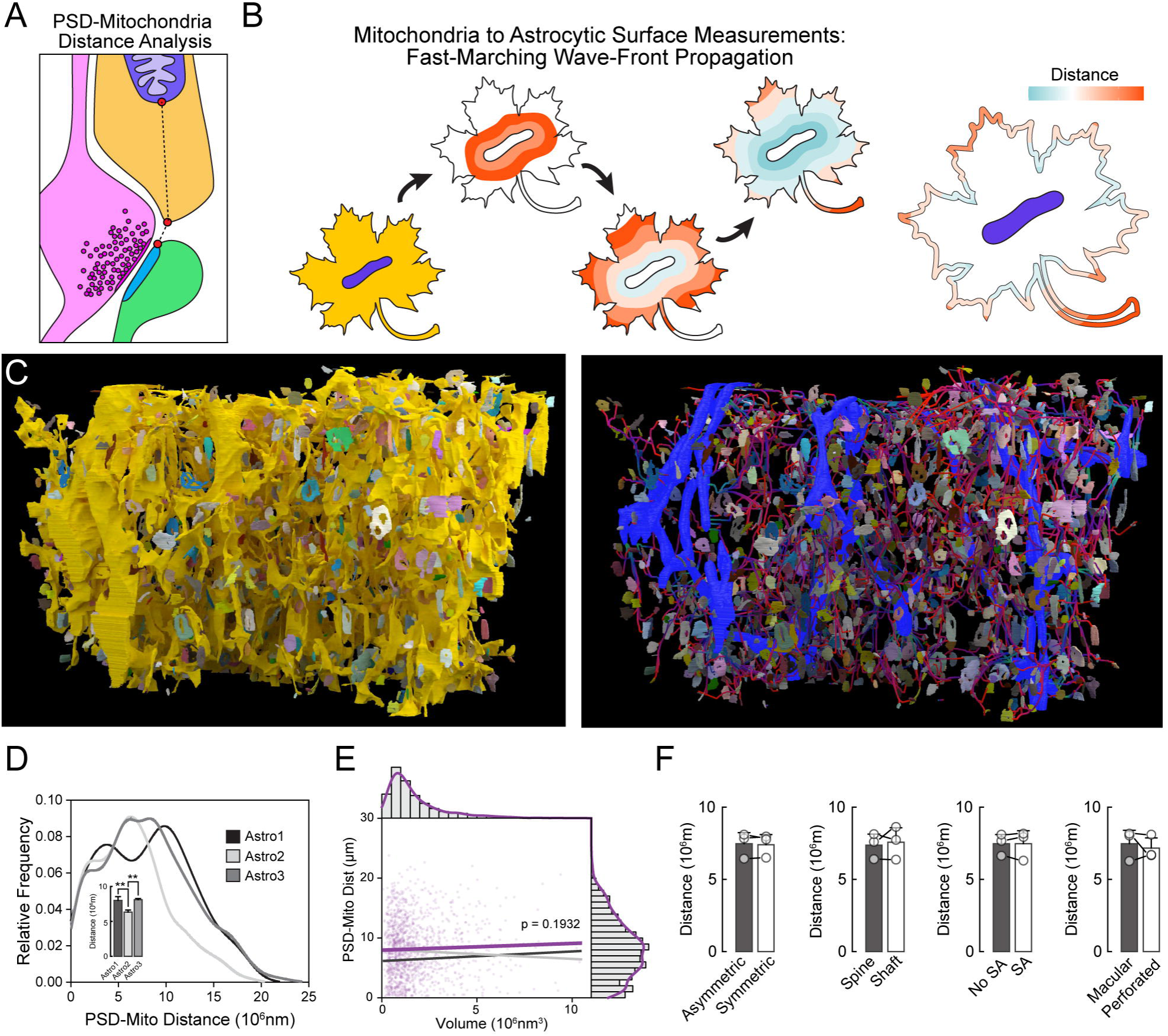
Mitochondria are not localized more closely to PSDs displaying particular characteristics. (A) Schematic showing geodesic measurements from PSD to astrocytic surface and astrocytic surface to mitochondrion. PSD to mitochondria distance is the sum of these two distances. (B) Fast-marching wave-front propagation to measure mitochondria to astrocytic surface shortest paths. (C) (D) (E) Correlation of PSD to mitochondria distance with PSD volume for astrocytes (Pearson Product Moment Correlation). (F) Summary plots showing that there were no significant differences between the proximity of PSDs of certain classes to the nearest astrocytic mitochondrion. Scale cube = 1μm^3^

### Synapses Organize into Discrete Clusters Around Astrocytic Nanoarchitecture and Show Cluster-Specific Proximity to Astrocytic Mitochondria

The multi-modal distributions of PSD-Mito distances suggested that there may be distinct PSD populations that lay nearer or farther from mitochondria (Figure 5D). Furthermore, we observed qualitatively that PSDs appeared to form distinct groups around certain astrocytic parts that are separated from the rest of the astrocyte by constrictions (Figure 6A-B). We therefore hypothesized that astrocytes have a specialized arrangement with groups of synapses, or “synaptic clusters.” Given the lack of an established “ground truth” test set for such synaptic clusters, and also given the high degree of 3D complexity of the astrocyte and PSDs surrounding it, a parameter-free, unsupervised clustering approach was required to decipher the relationship of PSDs to astrocytes. To accomplish this, we employed dominant sets (DSet) clustering, which has been used for a wide range of clustering problems, and requires no input other than a similarity matrix (Pavan and Pelillo, 2003b, 2003a, 2006). Using an eikonal flow-based approach, we calculated PSD-to-PSD pairwise distances to obtain complete similarity matrices for each astrocyte and applied a customized DSet clustering algorithm to these matrices (Figure 6C). We investigated PSD-PSD distances measured within the boundaries of the astrocyte (“intravolume distance;” Figure 6C, Figure 7) as well as distances measured along the surface of the astrocyte (“surface distance;” not shown). Remarkably, we found that synapses around astrocytes organize into discrete clusters that range in size from 4 to 55 PSDs per cluster (Figure 6D; Figure 7A). To ascertain whether astrocyte morphology did indeed play the operant role in the structure of these clusters, we performed DSet clustering on Euclidian PSD-PSD distance matrices as a negative control. These distance matrices were based on distances between PSDs in empty space, neglecting the presence of the astrocyte. This yielded clusters very different from the intra-volume and surface distance matrices (not shown). This is expected due to the convoluted morphology of the astrocyte volume and surface. Thus, synapses form clusters around astrocytes, suggesting that astrocytes partition the synaptic space such that synapses are related to one another relative to how astrocytes interact with them, in addition to where they fall in Euclidian space and within a neural circuit.

**Figure 6.**
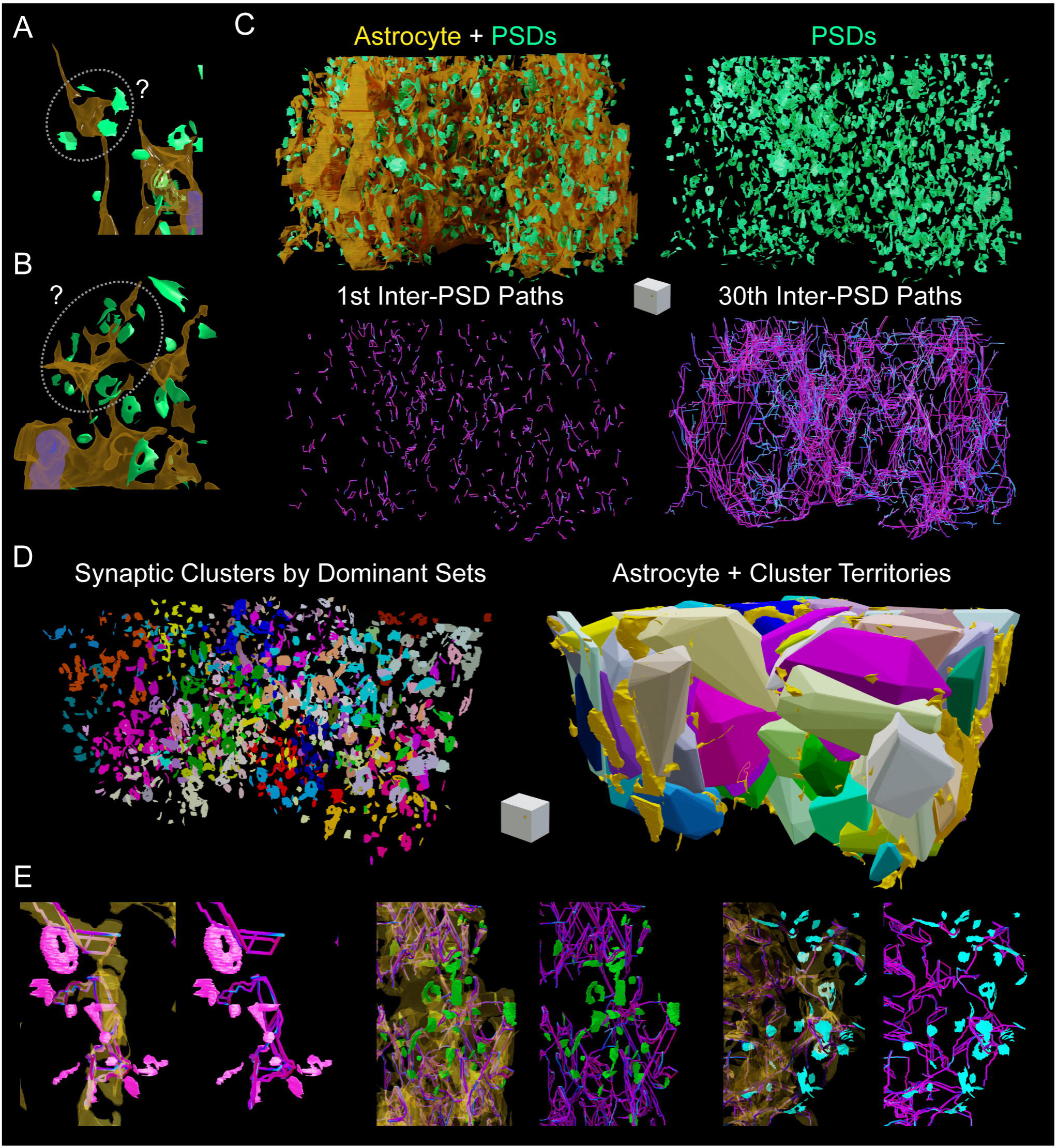
Investigating PSD organization around astrocytes and the identification of astrocyte-associated synaptic clusters. (A-B) Examples of astrocytic processes (yellow) interacting with PSDs (green). Dotted ellipses show apparent organization of synapses. (C) Geodesic measurements between PSDs. Upper Left, organization of all PSDs (green) around the astrocyte (yellow). Upper Right, display of all astrocyte-associated PSDs. Lower Left, example inter-PSD measurements (short paths; first nearest neighbors). Lower right, example inter-PSD measurements (long paths; 30^th^ nearest neighbors). (D) Analysis of inter-PSD measurements using undersupervised Dominant Sets clustering identifies synaptic clusters interacting with the astrocyte (left). Individual Dominant Sets clusters depicted as astrocytic-synaptic hubs. (E) Example Inter-PSD path measurements. Scale cube = 1μm^3^

**Figure 7.**
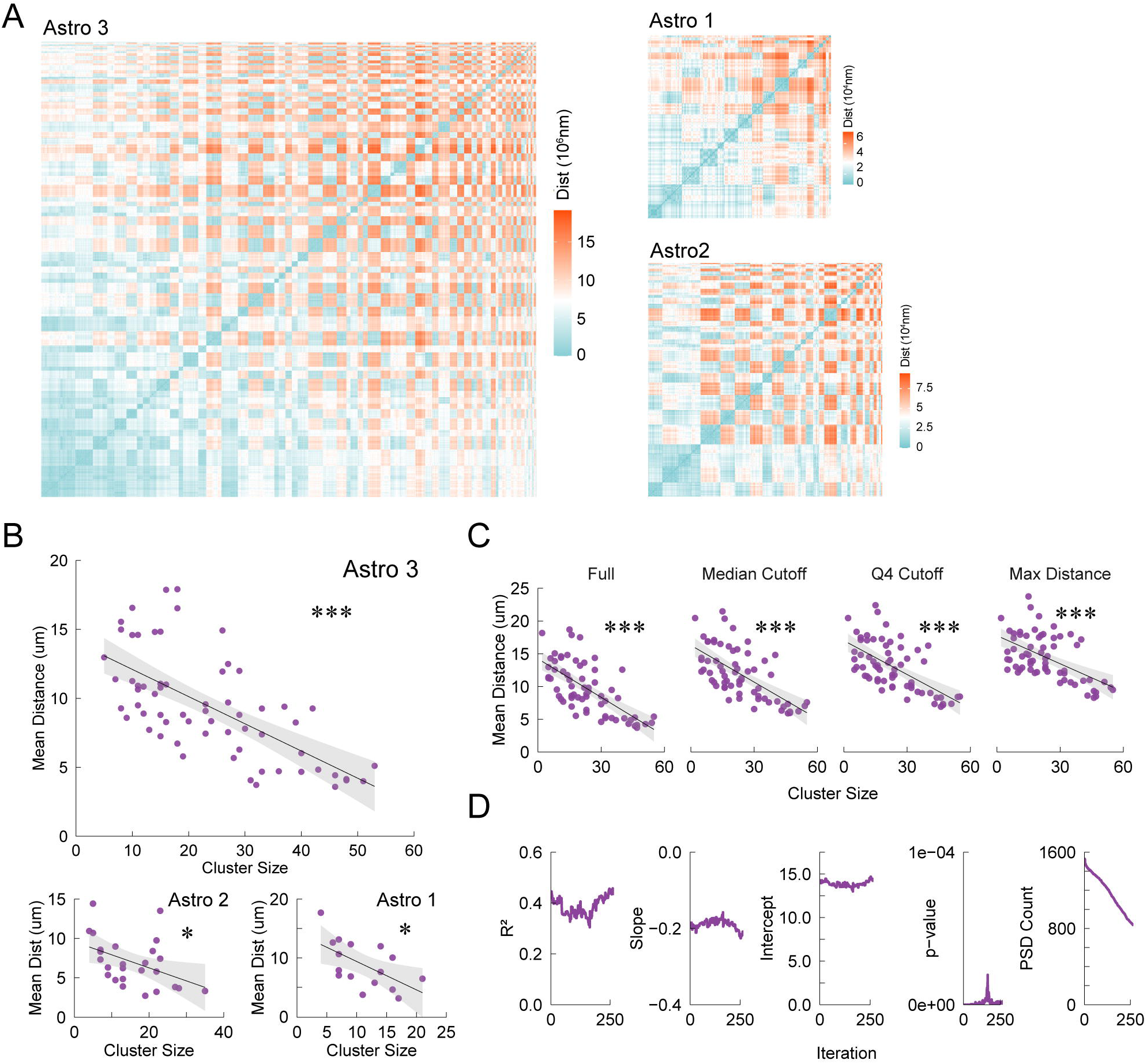
Astrocytes cluster analysis and proximity analysis of mitochondria to synaptic clusters. (A) Correlation matrices of distances between all pairs of PSD landing points along the surface of astrocytes in three reference volumes. The heatmap shows the distance between clusters. (B) Mean distance measurements showing that larger synaptic clusters are positioned closer to astrocytic mitochondria for Astro 3, 2, and 1. (C) Correlation of mean cluster PSD-Mito distance with cluster size was not influenced by removing the shortest 50%, 75%, or all but the longest distances from the mean cluster distances. Cluster size was not altered. (D) When the population of PSDs was incrementally eroded from the outside of the volume inward, by shrinking the bounding box for PSD inclusion while maintaining the astrocyte and mitochondrial surfaces as they are, mean PSD-Mito distances for the newly defined clusters still correlate strongly with cluster size, indicating that PSDs on the edge of the originally considered volumes do not impact the general organization of the clusters with respect to the astrocyte and its mitochondria. * p < 0.05, *** p < 0.001

We then asked whether the number of PSDs per cluster (cluster size), correlated with the proximity of clusters to their nearest mitochondria. When clusters were defined using either intra-volume or surface PSD-PSD distances, cluster size was inversely correlated with the mean distance of mitochondria to the PSDs in that cluster for all astrocytes (Figure 7B). This suggests that astrocytic mitochondria are positioned closer to large groups of synapses. As expected, PSD clusters based on the Euclidean PSD-PSD distance (i.e. neglecting the astrocyte), did not show a similar correlation (not shown). The robustness of this interaction between PSD-Mito distances and size of synaptic clusters was also demonstrated by removing the shortest PSD-Mito distances from the analysis (Figure 7C). Here, we repeated the correlation analysis for clusters that contained only the top 50%, top 25%, or the single longest PSD-Mito distance(s) for each cluster. This tested whether the observed correlation is simply driven by the fact that larger clusters might be expected to be closer on average to mitochondria by being spread out over more space. Even when only considering the longest distances, however, the correlation between cluster mean PSD-Mito distance and cluster size was present. We also asked whether edge effects may be present; PSDs close to the edge of the reference volume may actually be very near to a mitochondrion that is just outside the dataset. To account for this, we progressively removed PSDs from the edges of the dataset, performing a new round of clustering and regression each time a PSD was removed. As the population of PSDs considered was eroded, the summary data for the resulting cluster mean PSD-Mito distance versus cluster size correlation remained significant and otherwise relatively consistent (Figure 7D). This indicates consistency in the relationship between synaptic clusters and PSD-Mito distances and that PSDs on the edge of the volumes do not alter the relationship. Thus, astrocytes form higher-order interactions with clusters of synapses (rather than with individual synapses), and synaptic clusters predict the location of astrocytic mitochondria.

## DISCUSSION

Astrocytes are made up of a convoluted network of thin, irregular processes, that lack a readily apparent pattern or organization. Their myriad thin processes make intimate contacts with neurons to form the tripartite synapse, where they play key roles in regulating neural function (Perea et al., 2009). They also extend large processes that terminate in endfeet that surround vasculature and contribute to the maintenance of the BBB (Guérit et al., 2021). To understand the nanoarchitecture of astrocytes, we produced a series of high-resolution 3D volumes of astrocytes from mouse S1 cortex, and developed principled computational approaches to dissect their complexity. To our knowledge, no other studies that have utilized multiple serial EM datasets at equivalent resolution and scale to study astrocytic nanoarchitecture. Shape analysis of multiple volumes of astrocytes using computer vision techniques enabled their decomposition and cataloguing into 3 main part types including cores, expansions, and constrictions. Surprisingly, the majority of astrocytic parts end in terminations/tips, and rarely fold back onto themselves to form loops. Thus, astrocytes more closely resemble thorny bushes or shrubs with shallow, but wide networks of branches, rather than sponges with an abundance of looping structures and holes. We also found interesting structural features of astrocytic mitochondria, ER, and MERCs. Most prominent was the architecture of endfoot mitochondria and their convoluted tethering to the ER network through MERCs. Lastly, parameter-free cluster analysis revealed that astrocytic processes partition the neuropil and interact with discrete clusters of PSDs. Distance measurements and associated paths that accurately traverse convoluted 3D astrocytic morphology were made possible by a fast-marching method solution to the eikonal equation. The resulting measurements indicated that astrocytic mitochondria are positioned closer to large clusters of synapses. Altogether, these results demonstrate that astrocytes have specific geometry and nanostructural adaptations in the CNS.

The fine resolution of our datasets allowed a detailed analysis of mitochondria structure in the neuropil and endfoot regions. This showed that mitochondria shape and size vary significantly between subcompartments. The specific size and curvature of endfoot mitochondria was especially striking, potentially allowing these organelles to meet unique metabolic demands in this specialized area of the astrocyte. Indeed, mitochondria are dynamic organelles, constantly undergoing cycles of fission and fusion, which are important for mitochondrial quality control and for dealing with oxidative damage as well as for regulating mitochondrial morphology (Cagalinec et al., 2013; Sebastián and Zorzano, 2018). The shape of mitochondria is tightly linked to their bioenergetic properties and mitochondrial fission and fusion are regulated by metabolic signals. Adjustments of their morphology are thought to allow mitochondria to balance the rates of ATP production and metabolism of energy substrates according to local energy demands and nutrient supply, respectively (Liesa and Shirihai, 2013b; Mishra and Chan, 2016; Sebastián and Zorzano, 2018; Youle and Van Der Bliek, 2012). Mitochondrial hyper-fusion, resulting in highly elongated mitochondria is thought to optimize metabolic efficiency under conditions of nutrient starvation or metabolic stress (Tondera et al., 2009). Fragmentation, decreased mitochondrial length, and increased mitochondrial diameter on the other hand, has been linked to an excess of energy substrates. This conformation is thought to minimize the production of reactive oxygen species (ROS) through controlled metabolic uncoupling, in order to protect mitochondria from oxidative damage under conditions of excessive nutrient supply (Liesa and Shirihai, 2013a; Mishra and Chan, 2016; Sebastián and Zorzano, 2018; Youle and Van Der Bliek, 2012). This might be particularly important in astrocytic endfeet, given their proximity to the blood stream and their function as the primary sites of glucose uptake in the brain (Bélanger et al., 2011; Morgello et al., 1995). Ca^2+^ cycling between ER and mitochondria at MERCs is also known to regulate the metabolic capacity of mitochondria, by increasing mitochondrial matrix Ca^2+^ concentrations, which promotes oxidative phosphorylation (Cárdenas et al., 2010; Marchi et al., 2017; Raturi and Simmen, 2013; Simmen et al., 2010). The extensive MERCs observed in endfeet may allow for enhanced metabolic interaction of the two organelles and generate needed energy for supporting continuous transport across the BBB, as well as, for local protein synthesis (Boulay et al., 2017). Furthermore, their specialized structural properties may allow the ability to better replenish intracellular ATP stores, which has been shown to drop sharply after EAAT2/GLT-1 dependent glutamate uptake (Langer et al., 2017).

Protoplasmic astrocytes form mutually exclusive territories with neighboring astrocytes and have thousands of tiny processes that interact with pre- and postsynaptic sites through PAPs. We identified and analyzed thousands of synapses contacted by individual astrocytic PAPs across three reference volumes. Interestingly, it has also been previously observed that mitochondria were excluded from PAPs, with a minimal distance of ∼0.5μm between PSDs and any astrocytic Ca^2+^ source (i.e. mitochondrion or ER) (Patrushev et al., 2013). However, we found that in S1 cortex there is an appreciable portion of synapses within less than 0.5μm of mitochondria when considering PSD-Astro-Mito distance (Figure 5). Thus, mitochondria are not fully excluded from PAPs as previously thought. However, the PSDs that are closer to mitochondria do not appear to be of any particular type.

Unsupervised clustering was used to relate the location of PSDs with respect to the astrocyte surface. This revealed specialized domains in astrocytes created by higher-order clusters of synapses associated with astrocytic mitochondria. We found that there is a decrease in average distance to mitochondria with increasing cluster size, suggesting that the mitochondria that are closest to PSDs serve a large number of them. Moreover, this suggests there is an optimal organization of mitochondria within astrocytes to support metabolic load related to nearby synaptic clusters. Whether or not astrocytes help organize the clusters or astrocytes structurally adapt to synaptic clusters remains to be tested and will require developmental studies, chronic *in vivo* imaging, and new mouse genetic lines that can simultaneously tag specific astrocyte structural parts and astrocytic mitochondria to monitor their mobility. This can test if astrocytic mitochondria adjust their positioning, for example, through activity-dependent localization mechanisms (Jackson and Robinson, 2018).

Despite the significant advances in understanding astrocytic geometry and nanoarchitecture provided here, there are several technical limitations that warrant discussion and consideration in future studies. Here, we have presented subregions of astrocytes and therefore were not able to apply our computational analysis to entire cells. While conventional FIB-SEM generates high-resolution ultrastructural volumes (< 10 nm isotropic voxels) with robust registration of serial images (Xu et al., 2017), its field of view and z-depth are limited to a scale similar to the datasets in the present study. On the other hand, serial block face SEM (Calì et al., 2019) or automated tape-collecting ultramicrotome (ATUM) systems (Kasthuri et al., 2015) allow for much larger pieces of tissues to be imaged. However, these methods are constrained by section thickness due to the requirement of manual cuts with a diamond knife (typically ∼30-150nm) which, in our astrocytic nanoarchitectural analysis, would underestimate the frequency and geometry of many astrocytic constrictions (Figure 2D). FIB-SEM ‘volume stitching’, whereby 20μm thick slabs of embedded tissue are first collected with a conventional ultramicrotome and then imaged at mill depths of <10nm has shown promise in attaining z-resolution high enough to allow for accurate mapping of fine axons across gaps between slabs (Hayworth et al., 2015). Based on the current study <10nm mill depths within slabs should allow for adequate sampling of astrocyte morphology. It must, however, be noted that astrocytes can exceed 100 μm in diameter (Bushong et al., 2002). While the axial field of view accessible by advanced FIB-SEM techniques has improved enough to accommodate cells of this size (Hayworth et al., 2020; Xu et al., 2017), the ∼30nm gap size between tissue slabs in volume-stitched samples (Hayworth et al., 2015) may be problematic given the ubiquity of very fine processes in astrocytic volumes (Figures 2 and 3). Whether or not these approaches will allow for high quality reconstruction of fine astrocytic processes in larger volumes therefore remains to be seen.

Exceedingly thin astrocytic processes also present a problem for automated segmentation, introducing a major rate-limiting step for investigating astrocytic nanoarchitecture. Attempts were made in this study to utilize automatic or semi-automatic segmentation approaches (Berg et al., 2019; Cardona et al., 2012). However, the currently available methods were found to be inefficient, and failed to accurately trace astrocytic structure without time-consuming manual interventions. Instead, multiple rounds of manual segmentation and detailed quality control by experienced astrocyte biologists were required to find complete sets of connected astrocytic branches in our reference volumes (see Methods). Indeed, segmentation of nanoscopic branches of astrocytes are an acknowledged challenge in the field (Rusakov, 2015), and in fact, special attention must be paid to removing segmenting errors caused by the presence of astrocytes in current state-of-the-art automated segmentation approaches (Januszewski et al., 2018; Schubert et al., 2019). While crowdsourcing astrocytic segmentation followed by stringent quality control may be a viable alternative (Arganda-Carreras et al., 2015), astrocyte-specific auto-segmentation strategies are likely needed to reconstruct fine astrocytic nanostructure for complete cells and fully integrate astrocytes into the pursuit of saturated reconstructions and connectomes.

Regardless of the limitations listed above, we expect that the computational methods we have developed in this study will be useful in unravelling astrocytic nanoarchitecture in high resolution datasets regardless of the final approach used to achieve accurate, largescale models of whole astrocytes. Furthermore, our results can serve as a launching point for computational modeling of astrocytic function in the brain, using the models generated as ground truth descriptions and measurements of astrocytic geometry and complexity. Integrating these new datasets and analyses with biophysical modeling environments such as NEURON and simulation tools such as ASTRO (Savtchenko et al., 2018) is expected to improve our ability to comprehend astrocyte function. Finally, this study lays the groundwork for considering how astrocytes interact with surrounding microcircuitry at the highest levels of cellular resolution. Astrocytes and neurons are essential partners in nervous system function, and therefore unravelling their relationships to one another is be a central part of current and future connectomics work.

## MATERIALS AND METHODS

### Animals

Experiments were approved by the Montreal General Hospital Facility Animal Care Committee and followed the guidelines of the Canadian Council on Animal Care. Mice were housed and cared for according to standard operating procedures at McGill University and the Research Institute of the McGill University Health Centre.

### In Utero Electroporation

In utero electroporation (IUE) was performed as described (Matsui et al., 2011; Saito, 2006). Timed-pregnant female CD1 mice were purchased from Charles River Laboratories (Saint Constant, QC). Surgeries were performed on embryonic day 18. Pregnant mice were anesthetized using isoflurane, then an incision along the abdomen was made to expose the uterine horns containing embryos. After exposing the uterine horns, embryos were periodically flushed with warm (37°C) sterile PBS (Gibco, Thermo Fisher, Waltham, MA). Using pulled borosilicate pipettes that were bevelled with diamond lapping film (FOSCO, Fremont, CA), a mixture of 3 µl of DNA was injected into the lateral ventricles. Precise injection volume was attained with a foot pedal-controlled Elite Nanomite syringe pump (Harvard Apparatus, Holliston, MA) The DNA mixture contained 1 µg/µl of PB-GfaABC_1_D-Lck-mCherry to label the plasma membrane of astrocytes, 1 µg/µl of PB-GfaABC1D-mitoDendra2 to label mitochondria of astrocytes, and 1 µg/µl of the transposase-expressing PB200PA-1. To help visualize the injection of the DNA mixture, 0.1% (w/v) of Fast Green dye (Millipore Sigma, Oakville, Ontario) was added to the solution and adjusted to the final volume with sterile PBS. Following injection of the DNA, 3 mm tweezer-electrodes were used to apply 3 × 50 ms 42V pulses at 950 ms intervals via an ECM 830 square pulse generator (BTX, Holliston, MA). After injection of all the embryos, the uterine horns were placed back inside the abdomen and the incision site sutured. Pregnant mice were returned to their home cage and given post-surgical analgesics for 3 days and monitored until full recovery.

### DNA Constructs

Astrocytes and their precursors divide rapidly throughout postnatal development which makes episomal plasmid delivery less favorable due to the dilution of the DNA and low efficiency of gene expression following IUE. With the PiggyBac transposon system, DNA constructs are integrated into chromosomal DNA, thus resulting in stable gene expression (Chen and LoTurco, 2012; García-Marqués and López-Mascaraque, 2012). PiggyBac plasmids were purchased from SBI Systems Bioscience (Palo Alto, CA). All DNA constructs used for astrocyte targeted expression were manually cloned into the transposon plasmid (PB514B-2) under control of the modified human GFAP promoter, GfaABC_1_D (Lee et al., 2008) using restriction cloning and In-Fusion HD assembly (Clontech, Takara Bio, USA). The following sequence elements were obtained from Addgene: GfaABC_1_D (#19974) (Liu et al., 2008), Lck (#34924) (Akerboom et al., 2012), and mitoDendra2 (#55796) (Pham et al., 2012). The mCherry sequence was originally obtained from a plasmid donated by Dr. Roger Tsien (UCSD). DNA was prepared using the endotoxin-free NucleoBond Xtra Maxi EF kit (Clontech, Takara Bio, USA).

### Tissue preparation for Confocal Imaging and Structured Illumination Microscopy (SIM)

Electroporated mice (6 months old; male and female; CD1) were anesthetized with isoflurane and tissue fixed by transcardial perfusion first with ice cold DPBS followed by 4% paraformaldehyde in 0.1 M phosphate buffer (pH 7.4) at 5 ml/min for 5 minutes. Brains were removed and post-fixed in the same solution overnight at 4°C. Following equilibration in 30% sucrose in DPBS, the tissue was embedded in OCT (Somagen, Edmonton, AB) and snap frozen. Brains were cryosectioned coronally at 30 µm thickness and stored in DPBS at 4°C until use. Sections were mounted onto charged Superfrost™ Plus glass slides and preserved with SlowFade™ Gold Antifade Mountant (Thermo Fisher, Waltham, MA). Confocal microscopy was performed using an Olympus FV1000 laser scanning confocal microscope with a 60x oil-immersion objective (N.A. 1.4) at a resolution of 0.132 µm^2^/pixel. Stuctured Illumination Microscopy (SIM) image stacks were acquired using a Zeiss LSM880 ElyraPS1 microscope at the RI-MUHC Molecular Imaging Platform with a 63x oil-immersion objective (Plan-Apochromat 63x/1.40 Oil DIC). All images were obtained using 5 rotations and processed for SIM using the Zeiss Zen software. For both confocal and SIM, mCherry and mitoDendra2 were visualized using a 568 nm and 488 nm laser, respectively.

### Electron Microscopy Sample Preparation

Postnatal day 50 adult male C57/Bl6 mice were anaesthetized with ketamine-xylazine cocktail and transcardially perfused with 9-10 ml of 0.1M phosphate buffer (PB) at room temperature (RT) at 7 ml/min, followed by 30-40 minutes of perfusion with 2.5% glutaraldehyde, 2% formaldehyde, 0.1M PB at 7 ml/min. Brains were removed from the skull and allowed to post-fix in the same fixative solution at 4°C for 6-12 hours. Tissue chunks containing somatosensory (S1) cortex were manually dissected based on visible landmarks using straight razor blades and cut into 300 μm slices using a vibratome (Knott et al., 2011; Korogod et al., 2015). Tissue preparation, staining, and processing for FIB-SEM was performed based on established protocols by the Facility for Electron Microscopy Research of McGill University. Briefly, following postfixation, the tissue was washed 3 times in 0.1 M sodium cocodylate buffer and stained with 1.5% potassium ferrocyanide and 1% osmium tetroxide/0.1 M cocodylate for 30min. Tissue was then washed in ddH20 twice for 5 min, and block stained in 1% uranyl acetate in ddH20 for 30 min and washed twice with ddH20. Tissue was passed through an acetone/ddH20 dehydration series of 30%, 50%, 70%, 80%, 90% and 3x100% acetone for 2 min at each step. This was followed by an epon infiltration series of 1:1, 2:1 and 3:1 epon:acetone for 30 min at each step. Tissue was left in 100% epon for 6 hours before being allowed to harden at the point of a conical mold at 65°C for 24 hours. Thin sections were prepared and stained with toluidine blue to select the area for imaging. Ultrathin sections were examined by transmission electron microscopy on a Tecnai 12 BioTwin 120kV TEM equipped with an AMT XR80C CCD Camera (FEI, Hillsboro, OR) to assess tissue and staining quality.

### Focused Ion Beam Scanning Electron Microscopy (FIB-SEM)

FIB-SEM imaging was performed as described previously (Morita et al., 2017). Epon blocks containing tissue were trimmed with a razor blade to expose the region of interest, then mounted on a 45° pre-titled SEM stub and coated with a 4nm layer of platinum to enhance electrical conductivity. Gallium ion beam milling of serial sections and block face imaging after each mill were carried out on a Helios Nanolab 660 DualBeam system using Auto Slice & View G3 ver 1.5.3 software (FEI). The sample block was first imaged to determine the orientation of the block face and ion and electron beams. A protective platinum layer 80 μm long, 19 μm wide and 2 μm thick was deposited on the surface orthogonal to the block, where the ion beam is first incident, to protect the sample from ion beam damage and to correct for stage and/or specimen drift. Trenches were milled on both sides of the region of interest to minimize re-deposition of milled material during automated milling and imaging. Fiducial markers were generated for both ion and electron beam imaging and were used to dynamically correct for drift in the x- and y-directions during data collection by applying appropriate scanning EM beam shifts. Milling was carried out at 30 kV with an ion beam current of 9.3 nA, stage tilt of 6.5°, and working distance of 4 mm. At each step, an 8nm slice of the block face was removed by the ion beam. This mill depth was chosen to ensure resolution of the smallest astrocytic processes while still collecting images of an appreciable volume. Each newly milled block face was imaged simultaneously with the Through the Lens Detector (TLD) for backscattered electrons and In-Column Detector (ICD) at an accelerating voltage of 2 kV, beam current of 0.4 nA, stage tilt of 44.5°, and working distance of 3 mm. The pixel resolution was 4.13 nm with a dwell time of 30μs per pixel. Pixel dimensions of the recorded image were 3072 × 2048 pixels. The acquired image stacks measured 12.7 × 8.5 × 4.4 μm for Volume 1 (551 images), 12.7 × 8.5 × 5.4 μm for Volume 2 (672 images) and 12.7 × 8.5 × 7.8 μm (976 images).

### Stack Alignment, Segmentation, and Quality Control

Image stack alignment and manual reconstruction were performed using TrakEM2 (Cardona et al., 2012). Astrocytes were identified by clear cytoplasm devoid of visibly oriented microtubules, the presence of dark staining glycogen granules, and the absence of synaptic inputs (Spacek, 1985a and personal communication with Dr. Spacek). We also noted that in our preparations, astrocytic mitochondria stained more lightly than neuronal mitochondria. Due to the extremely convoluted structure of astrocytes and the fact that some branches decreased with only a couple pixels in width (corresponding to 4-8 nm), extensive quality control was required. Two to four rounds of quality control (QC) were carried out by BK and CKS (at least one round each) on each electron micrographs to ensure accuracy. Corrections were applied when necessary. To perform QC, we overlaid a grid onto the TrakEM2 workspace and panned through the entire volume within each square of the grid, flagging errors and omissions for later confirmation and segmentation. Generally, new PSDs and new astrocytic branches (sometimes quite large), were discovered during QC. Segmentation of thin astrocytic processes was stopped when astrocytic membranes could not be differentiated from one another. In a number of such cases, it did appear that the astrocytic membrane was simply pinched, rendering the process too thin to be resolved, thus a small number of processes may extend further than the segmented volume shows. All 3D models were generated using Blender unless otherwise specified. As mentioned, astrocytic filaments can be extremely thin. The reduced resolution provided by backscattered electrons meant that the interstitial space and occasionally the band of cytoplasm in protoplasmic processes could not be distinguished. In these cases, when it was unclear whether there was an obvious continuation of cytoplasm on either side of the restriction, the connection was not traced. Thus, we likely underestimated the extent of continuous astrocyte in the volumes considered.

### Astrocyte Shape Analysis

We characterized the local shape of the astrocytic surface by computing its principal curvatures, or equivalently, the Mean and Gaussian curvatures. A convenient way to do this is to use an outward Euclidean distance function to the astrocyte surface as an embedding, and then to use numerical measures that operate directly on this embedding as is often done for numerical methods for front propagation (Osher and Fedkiw, 2003; Sethian, 1999). For example, the Mean curvature at any point on the astrocytic surface is the divergence of the normalized gradient vector of this Euclidean distance function embedding. In addition to surface curvatures, we have employed a novel way to characterize local astrocytic width at any point on the surface. We do this by using twice the radius of the maximal inscribed sphere within the astrocytic volume, that touches that point (Siddiqi and Pizer, 2008), to define local thickness. Such a sphere has the property that it fits within the astrocyte volume while just touching it on at least two points. Thus, any larger sphere that contains such a sphere, and shares its center, must protrude outside the astrocytic volume. To compute this local width efficiently we use 3D average outward flux based medial surfaces (Siddiqi et al., 2002). We also partition the interior of the astrocytic volume into compartments by labeling each point in the volume by the identity of the mitochondrion it is closest to. In doing so we get an accurate view of how the shape of the mitochondria and their 3D arrangement within the astrocyte volume determine and influence the minimal paths to the astrocyte surface.

For astrocyte endfoot structure, blood vessels were classified as capillaries due to their small diameter (4-5µm), absence of juxtaposed smooth muscle cells, and the presence of a single thin layer of basement membrane (BM). Surface curvatures for reconstructed ER and mitochondria were calculated from triangulated surface meshes created in Blender. Blender generated surface meshes were smoothened to reduce segmentation artifacts by applying a combinatorial Laplacian operator over 3 iterations using the Graph Theory Toolbox (https://www.mathworks.com/matlabcentral/fileexchange/5355-toolbox-graph). Principal curvatures were calculated as described above. ER surface points were clustered based on their total curvature (|k1|+|k2|) using a Gaussian mixture model that was fit to the data using the fitgmdist function in MATLAB. A Gaussian mixture model with 3 components was fit to the combined total curvature data of all reconstructed ER segments and surface points were clustered based on this model. ER curvature calculations and clustering were performed on the GRAHAM (GP3) computing cluster at the University of Waterloo, through the Compute Canada (www.computecanada.ca) network.

### PSD Segmentation and Profiling

We segmented all PSDs contacted by segmented astrocytes in reference volumes. PSDs were segmented manually. PSDs were considered to be associated with the astrocyte if the perisynaptic astrocytic process (PAP) made contact anywhere on the synaptic unit (defined as the synaptic cleft, spine head, spine neck and axonal bouton)(Genoud et al., 2006). For shaft synapses, since the variable size of the postsynaptic dendrites meant that the postsynaptic compartment could be very large, only shaft synapses contacted on the axonal varicosity or at the synaptic cleft were considered. Only PSDs that had clear symmetric or asymmetric dark staining and had at least three apparent vesicles were segmented (Knott et al., 2002). Nearly all PSDs were associated with many docked and undocked vesicles. In each volume, a small proportion of PSDs were omitted from analyses as they were cut off at the edge of the volume and hence partial or distorted due to FIB-milling artifacts.

PSDs were categorized by: 1) their location (spine, dendritic shaft, cell body); 2) their type (asymmetric or symmetric); 3) the presence or absence of a spine apparatus (SA); 4) whether they were macular or perforated; 5) whether they were associated with endoplasmic reticulum (ER) at the PSD, spine base or in the spine neck; 6) whether they were associated with a mitochondrion that directly overlapped in 3D with the PSD or the spine base; and 7) whether the synapse-associated ER formed a MERC with a mitochondrion somewhere in the postsynaptic neuron. It is important to note that mitochondrial and MERC distance categories are complicated by the fact that we considered only small proportions of total astrocytes, and the astrocytes extend outside the volumes we imaged, meaning that mitochondria or MERCs could occur just out of frame. This was mitigated in two ways: PSDs that were close to the edge of the block were not categorized for their association with mitochondria or MAMs, and only a minority of synapses where on dendrites that completely lacked mitochondria or MAMs.

### Dominant Sets Clustering of PSDs

Most clustering methods, such as k-means, seek a strict partition of the entities into disjoint sets. We chose instead to use the dominant sets (Dsets) clustering algorithm (Bulò and Bomze, 2011; Pavan and Pelillo, 2006; Rota Bulò and Pelillo, 2017) which provides a formal definition of a cluster for each set of PSDs grouped and allows for a clear theoretical interpretation of the results. This method requires as input a measure of pairwise similarity between each pair of entities. These similarities are expressed in terms of the entries in an adjacency matrix. The method then finds clusters with two guarantees. First, entities (PSDs) within the same cluster have a high degree of internal coherence, i.e., there is a high degree of similarity between any pair of PSDs within the same cluster. Second, the clusters are locally maximal, in that no cluster can be absorbed within a larger one, without decreasing the average pairwise similarities.

Our application of the Dsets algorithm used a similarity score that is inversely proportional to the geodesic distances between pairs of PSDs either within the astrocyte volume (intra-volume distance) or along the surface of the astrocytes (surface distance). We associated each PSD to the closest point on the astrocyte surface. Geodesic distance was then computed between points on astrocyte surface closest to each PSD using fast marching. To compute pairwise distance from each PSD surface point to every other PSD surface point on the astrocyte, we repeated the process with every PSD as the starting point. We experimented with several monotonically decreasing functions of geodesic distance in constructing the similarity weights of the adjacency matrix, all of which gave essentially the same results.

In addition to its theoretical guarantee that each cluster is a dominant set, the algorithm provides additional advantages for clustering PSDs. The number of clusters is derived from the data itself – there is no requirement that this be specified a priori. The approach allows for PSDs to belong to more than one cluster, which can be helpful for allowing for partial overlap in the PSD assignments, when appropriate. Finally, for our case of symmetric affinities, the solutions are maximizers of a quadratic function and can be efficiently found by using an appropriate dynamical system. In practice, both replicator dynamics and infection and immunization dynamics (Bulò and Bomze, 2011; Rota Bulò and Pelillo, 2017) give the same results while the latter is more efficient for large problem sizes.

### PSD-to-Mitochondria Distance Measurements

We used a fast marching method (Sethian, 1996) to solve the eikonal equation (Arnold, 1978) both inside and outside the segmented astrocytes to compute distance from mitochondria to PSDs. The segmented mitochondrial surfaces were used as the initial wave-front inside the astrocytic volume, and the astrocytic surface was used as the initial wave-front outside the astrocytic volume. The fast marching approach allowed us to evolve the initial surfaces to compute distance from mitochondria at every point inside the astrocyte volume and from the astrocyte to every point outside the astrocyte. We assumed a constant isotropic metric everywhere inside and outside the astrocytic surface. The total distance to each PSD was taken as the sum of the distance from the PSD to astrocyte surface and the distance from mitochondria to the closest astrocytic surface point of the PSD. To compute shortest paths, we use gradient descent on the computed distance map.

(Arnold, 1978)(Sethian, 1999)This computation is accurate, while also considering the complex 3D arrangement and shapes of all mitochondria within the astrocyte, together with the convoluted shape of the astrocyte itself, and the distribution of PSD regions outside it. This provides a powerful and significant advantage over the alternatives of using Euclidean distance, and heuristic measures of local surface to volume ratios for characterizing PSD distances.

### Statistical Analysis

Statistical analyses and graph plotting were performed using Origin (OriginLab, Northampton MA), Matlab (MathWorks, Natick MA) and R (R Core Team, 2021). R packages used included ggplot2 (Wickham, 2016), lemon (Edwards, 2020), scales (Wickham and Seidel, 2020), dplyr (Wickham et al., 2021), tidyr (Wickham, 2021), igraph (Csardi and Nepusz, 2006), and dbscan (Hahsler et al., 2019). Unless otherwise noted, mean comparisons were performed using Student’s t-test or ANOVA and 2-Way ANOVA with Tukey HSD pairwise comparisons, preceded by the Shapiro-Wilk normality test.

For mitochondria/ER/MERC analysis, where conditions of normality were met, we performed Welch’s unequal variances t-test, otherwise the non-parametric Mann-Whitney test to compare experimental groups. For comparing the size of individual manually segmented MERCs, data was log_10_ transformed to satisfy the conditions of normality for Welch’s unequal variances t-test. We calculated the mean and 95% confidence interval of log transformed data. The reported values are the anti-log of the mean +/- confidence interval. The anti-log of the log mean corresponds to the geometric mean of the population.

## ACKNOWLEDGEMENTS

We would like to acknowledge members of Murai and Siddiqi labs for their input on the experimental set-up and computational analysis applied in this study, as well as Dr. Donald van Meyel and Dr. Arjun Krishnaswamy for helpful comments on the text. Many thanks to Dr. Kelly Sears, Jeannie Mui, and Dr. Weawkamol Leelapornpisit at McGill’s Facility for Electron Microscopy Research (FEMR) for technical support with EM sample preparation and FIB-SEM. We are also particularly indebted to Dr. Tyler Sloan of Quorumetrix Solutions for extensive work and advice with regards to 3D models and Blender.

## AUTHOR CONTRIBUTIONS

KKM, KS, CKS, JBK, and TS conceived the project, analyzed data, and wrote the manuscript. CKS, JBK, and AS performed wet-lab experiments. TS, CKS, JBK, KS and KKM designed and/or implemented computational approaches for quantitative structural analysis and TS, CKS, and JBK produced the code for analysis. NA, MP, MPR, MG, AS, BK and CKS contributed to segmentation, reconstruction, and visualization of astrocytes. NA contributed to quantitative analysis of astrocyte endfeet, mitochondria and ER. HV and CAM contributed to the early conception of the project and discussions of adaptation and implementation of FIB-SEM technology.

---

**Supplemental Figure 1.**
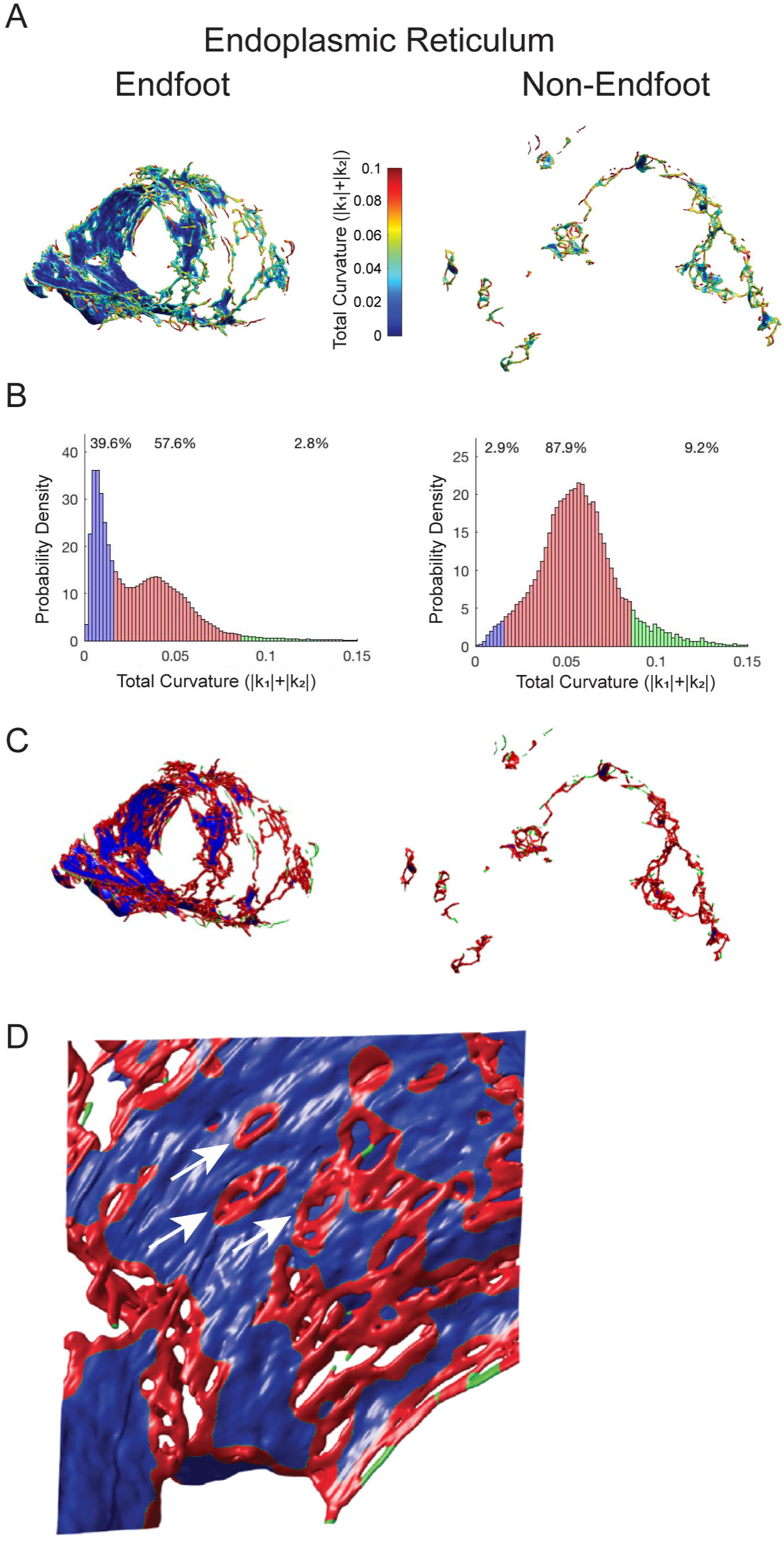
Nanostructural characterization of ER surface in astrocytic endfeet. (A) Example reconstructed endoplasmic reticulum from a reference volume in endfoot (left) and non-endfoot regions (right). Surface colour indicates total curvature (|k1|+|k2). (B) Points on the ER surface were clustered based on surface curvature to quantify the percentages of the ER using a Gaussian mixture model (GMM). This reveals sheet-like cisterns (blue), tubules (red), and small extensions (green) of ER. (C) Reconstructed ER with surface colours corresponding to GMM clustering results. (D) Representative example of fenestrations (arrows) in large flat cisterns in endfeet ER.

